# Gut-derived GLP-1 released by rare sugar D-allulose cooperates with insulin to activate left-sided vagal afferents and enhance insulin sensitivity

**DOI:** 10.64898/2026.01.09.697679

**Authors:** Kento Ohbayashi, Mamoru Tanida, Chikara Abe, Hirotaka Ishihara, Wataru Omi, Naoto Kubota, Daniel J. Drucker, Toshihiko Yada, Yusaku Iwasaki

## Abstract

Glucagon-like peptide-1 receptor agonists (GLP-1RAs) ameliorate hyperglycemia by directly stimulating insulin secretion from the pancreas. In contrast, the physiological role of short-lived endogenous GLP-1 remains unclear, largely because of its limited access to pancreatic β cells. Here, we show that D-allulose-induced intestinal GLP-1 secretion (AIGS) cooperates with insulin to reduce blood glucose levels by enhancing insulin action, rather than insulin secretion, in male mice. This cooperation and remote signaling require left-sided vagal afferents forming the common hepatic branch, but not right-sided afferents. AIGS-enhanced insulin action required both GLP-1 receptors and insulin receptor substrate 2 in these neurons. Remarkably, AIGS improved insulin resistance and hyperglycemia more rapidly and potently than the GLP-1RA exendin-4. These findings reveal that a subclass of vagal afferent neurons synergistically activated by endogenous intestinal GLP-1 and insulin does not stimulate insulin secretion but augments insulin action to improve glucose tolerance. This novel extra-pancreatic GLP-1 action mediated by vagal afferents provides a promising basis for innovative type 2 diabetes therapies.

**ARTICLE HIGHLIGHT:**

- Compared with GLP-1 receptor agonists, the physiological roles and mechanisms of endogenous, short-lived GLP-1 in glucose metabolism remain poorly understood.
- We utilized the rare sugar D-allulose, a noncaloric GLP-1 secretagogue, as a tool to elucidate the physiological actions of endogenous GLP-1.
- D-allulose–induced intestinal GLP-1 release cooperates with insulin to activate left-sided vagal afferents, enhancing insulin action rather than insulin secretion and thereby regulating glycemic control.
- Because this acute mechanism improved hyperglycemia in type 2 diabetes more effectively than GLP-1 receptor agonists, targeting GLP-1/insulin–vagal signaling may inform novel therapies and dietary or nutritional interventions for T2DM.

## INTRODUCTION

Type 2 diabetes mellitus (T2DM) and obesity have emerged as global health crises (1,2). A key pathophysiological feature of both conditions is insulin resistance, defined as impaired insulin responsiveness in major metabolic tissues such as skeletal muscle and liver (3,4). Preventing and reversing insulin resistance is essential for the effective management of T2DM. However, no established therapies currently offer both efficacy and safety in improving insulin resistance.

Glucagon-like peptide-1 receptor agonists (GLP-1RAs) with an expanded half-life in the circulation directly act on pancreatic β-cells to enhance glucose-induced insulin secretion and effectively ameliorate hyperglycemia. They are employed as drugs to treat T2DM (5,6) and recently obesity (7,8). In contrast, endogenous intestinal GLP-1, released in response to meal stimuli, has a short half-life of 1-2 min due to renal clearance and rapid degradation by dipeptidyl peptidase-4 (DPP-4) (9,10). Endogenous GLP-1 regulates glucose homeostasis independently of the GLP-1R expressed in pancreatic β-cells (11), suggesting extra-pancreatic targets. The physiological functions of endogenous GLP-1 and underlying mechanisms remain incompletely understood (12).

Recent studies have demonstrated that intestinal GLP-1 increases insulin secretion and regulates postprandial hyperglycemia through neural input using vagal afferent nerves (13–16). We have recently reported that intestinal GLP-1 release induced by the non-caloric stimuli of rare sugar D-allulose or gastrointestinal distention activates vagal afferent nerves, thereby enhancing insulin action and improving glucose tolerance (17,18). Notably, in high-fat diet-induced obese mice, D-allulose-induced GLP-1 secretion rapidly and significantly ameliorated hyperglycemia via GLP-1R signaling without stimulating insulin secretion, suggesting an action to improve systemic insulin sensitivity (17). However, this action remains to be established and the underlying mechanism to be clarified.

In this study, we investigated the role of D-allulose-stimulated GLP-1 secretion for insulin action in male mouse models of healthy, T2DM, and T1DM. We found that D-allulose markedly improved hyperglycemia in hyperinsulinemic T2DM mice by enhancing insulin action without stimulating insulin secretion, whereas it showed no effect in T1DM mice. We further found that D-allulose-induced intestinal GLP-1 acts in concert with insulin above basal levels to robustly activate vagal sensory pathways, particularly those mediated by GLP-1R and IRS2 in the left common hepatic branch, thereby increasing systemic insulin sensitivity. Moreover, while D-allulose–stimulated GLP-1 release did not lower blood glucose in healthy mice, it improved hyperglycemia more efficaciously than GLP-1 receptor agonists in T2DM mice with severe insulin resistance. Taken together, our findings reveal that intestinal GLP-1 and sufficient insulin cooperate to activate left vagal sensory neurons and enhance insulin sensitivity, which may provide a safe and effective strategy to improve insulin resistance in T2DM.

## RESEARCH DESIGN AND METHODS

### Animals

All experiments were approved by the Institutional Animal Experiment Committee of the Kyoto Prefectural University and in accordance with the Institutional Regulations for Animal Experiments (approval numbers: KPU060327-RC-3, KPU060327-RC-4, KPU060327-RC-5, KPU060327-C1, and KPU060327-C4).

Male C57BL/6J and db/db mice were purchased from Jackson Laboratory Japan, and Akita mice (Ins2 mutation) (19,20) were obtained from Japan SLC. Glp1r knockout (21) and Irs2flox/flox mice (22) were kindly provided by Dr. Daniel J. Drucker and Dr. Naoto Kubota, respectively. Glp1r-ires-Cre (23) and Phox2b-Cre mice (24) were obtained from Jackson Laboratory (strain no. 029283, 016223).

Mice were individually housed under controlled temperature (22.5 ± 2°C) and humidity (55 ± 10 %) with a 12-hour light/dark cycle (lights on at 7:30 am). Diet-induced obese mice (35–60 g) were produced by feeding a high-fat diet (D12492; Research Diets) for 50–100 days. Adult male mice (8–30 weeks) were used for all behavioral experiments.

### Surgical vagotomy and chemical deafferentation

As previously described with slight modifications, surgical vagotomy of either the common hepatic branch or subdiaphragmatic dorsal trunk of the vagus nerve (17,25), and chemical deafferentation of the common hepatic branch (25–27) were performed under anesthesia with a three-drug anesthetic of medetomidine (0.75 mg/kg; Nippon Zenyaku Kogyo), midazolam (4.0 mg/kg; Maruishi Pharmaceutical), and butorphanol (5.0 mg/kg; Meiji Seika Pharma). After either procedure, atipamezole (0.75 mg/kg; Nippon Zenyaku Kogyo) was administered for anesthesia recovery. Behavioral experiments were conducted at least 1 week post-surgery.

### Systemic desensitization of capsaicin-sensitive sensory nerves

Systemic capsaicin treatment was performed to impair capsaicin-sensitive sensory nerves including vagal sensory nerves, as previously described (18,28). Behavioral experiments were conducted at least 1 week after final capsaicin treatment.

### Microinjection of AAV vector into the nodose ganglion

Microinjection of adeno-associated virus (AAV) into the nodose ganglion (NG) was performed with modifications to previously described methods (17,29). Glp1r-Cre mice were anesthetized with a three-drug anesthetic, and the NG was exposed and microinjected with AAV2-hSyn-DIO-hM4Di-mCherry (1.0 × 10¹² GC/ml; #44362, Addgene) containing0.05% Fast Green (1–2 µl). Atipamezole was administered to facilitate anesthesia recovery. Behavioral experiments were conducted at least 3 weeks post-surgery. Following the behavioral testing, immunohistochemistry was conducted to verify hM4Di-mCherry expression, using rabbit anti-DsRed (#632496, 1/500; TaKaRa Bio) and, goat anti-rabbit Alexa Fluor 594 (#A-11012, 1/500; Thermo Fisher Scientific) antibodies.

### Measurement of GLP-1 in portal vein plasma

Mice were fasted for 4 h (9:00–13:00) before oral gavage of D-allulose (1 g/10 ml/kg) or saline. Portal vein blood was collected 60 min after administration under isoflurane anesthesia. Plasma was separated (4,000 rpm, 10 min, 4°C), and total GLP-1 was measured by ELISA (EZGLP1T-36K; Millipore). GLP-1 receptor antagonist exendin(9–39) (600 nmol/5 ml/kg, ip; ab141101, Abcam) was given 15 min before D-allulose (17).

### Sequential blood glucose and plasma insulin measurement and glucose, insulin and glibenclamide tolerance test

Mice were fasted for 4 h or overnight (18:00–10:00 the next day). Sequential measurements, baseline blood were collected from the tail vein, and D-allulose (1 g/kg) was administered either orally or intraperitoneally, or exendin-4 (0.5 nmol/10 ml/kg; ab120214, Abcam) was ip administered. Blood glucose was determined using GlucoCard Plus Care (Arkray) or Wako Autokit glucose C2 (FUJIFILM Wako) when over 600 mg/dl. Plasma insulin were measured by ELISA (MS303, Morinaga). For glucose, insulin and glibenclamide tolerance test, baseline blood glucose was measured, followed by oral gavage of saline or D-allulose (1 g/kg), and at 60 min post-administration, glucose (1 or 2 g/kg), insulin (1 or 1.5 IU/kg), or glibenclamide (1.5 mg/kg; 0.1% DMSO, 2% tween 80, 97.9% saline) was ip injected. Clozapine *N*-oxide (CNO; 3 mg/10 ml/kg, ip) was administered 1 h before gavage D-allulose to induce chemogenetic inhibition. Exendin(9–39) at 600 nmol/kg was injected ip 15 min before D-allulose or insulin administration.

### Measurement of food intake

Individually housed mice were fasted for 3 h before the dark phase with water available. D-allulose (1 g/kg) or saline was administered orally 10 min before lights-off, and food intake was measured.

### Immunohistochemical detection of pERK1/2 in nodose ganglion

Mice fasted for 4 h received oral saline or D-allulose (1 g/kg), followed 15 min later by intraperitoneal vehicle or glibenclamide (1.5 mg/kg). After another 15 min, mice were transcranial perfused with 4% paraformaldehyde solution. NGs were stained with anti-pERK1/2 antibody (#9101, 1:500; Cell Signaling Technology) and Alexa 488 goat anti-rabbit IgG (#A-11008, 1:500; Thermo Fisher Scientific). The number of pERK1/2-positive NG neurons in four sections per mouse was counted and averaged.

### Measurement of hepatic vagal sensory nerve activity

Mice were anesthetized with 2.0% isoflurane delivered through a polyethylene tracheal cannula connected to a respirator, and rectal temperature was maintained at 35.5 ± 0.5°C. Polyethylene catheters were inserted into the carotid artery and jugular vein for continuous blood pressure monitoring and drug injection, respectively.

The activity of vagal sensory nerve of the common hepatic branch was recorded as described in our previous study (27). The central side of nerve projecting to the liver was transected, and the peripheral segment was hooked onto bipolar stainless-steel electrodes, lifted and covered with silicone gel. Nerve signal was amplified and filtered using an amplifier, visualized on an oscilloscope, converted to a digital signal by an analog-to-digital converter, and recorded on a computer. After a stable recording period was confirmed, saline or D-allulose (1 g/10 ml/kg) was administered intragastrically via a polyethylene catheter, followed 60 min later by intravenous injection of vehicle or glibenclamide (1.5 mg/10 ml/kg).

After euthanasia, electrical noise was recorded to detect pure neural activity as spikes per 5 seconds. The average number of spikes per 1 min before intragastric injection was defined as 100%, and the change after injection was expressed as a percentage of baseline.

### Quantitative PCR analysis

Total RNA was extracted using either RNAiso Plus (TaKaRa Bio) or a combination of ISOGEN II and Etachinmate (Nippon Gene), and cDNA was synthesized using ReverTra Ace with an included DNase treatment (Toyobo). Real-time PCR was conducted with THUNDERBIRD Next SYBR qPCR Mix (Toyobo) on a CFX Connect system (Bio-Rad). Gene expression was quantified by the ΔΔCt method and normalized to 36b4. Primer sequences are listed in Supplementary Methods.

### Quantification and statistical analysis

Data are presented as means ± SEM. Statistical analyses were performed using two-tailed unpaired t-tests or one-way/two-way ANOVA with appropriate post hoc tests (Dunnett’s, Tukey’s, or Bonferroni’s). Analyses were conducted using Prism 10 (GraphPad Software). P < 0.05 was considered statistically significant.

### Data availability

All data supporting this study are included in the main text or the supplemental materials. Additional information is available from the lead contact upon request.

## RESULTS

### D-Allulose-induced gut GLP-1 release enhances insulin action and lowers blood glucose exclusively in type 2 diabetic mice via GLP-1 receptor activation

A single oral gavage of D-allulose significantly increased total GLP-1 concentrations in portal vein plasma in healthy C57BL/6J mice, T2DM *db/db* mice, and T1DM Akita mice. (Fig. 1A, 1D, 1G, 1J), without altering glucagon levels (Supplementary Fig. 1). Despite, robust increases in gut-derived GLP-1, blood glucose and plasma insulin levels remained unchanged following D-allulose administration in healthy mice, Akita mice, and streptozotocin-treated mice lacking circulating insulin (Fig. 1B–C, 1E–F and Supplementary Fig. 2). In contrast, hyperinsulinemic T2DM *db/db* mice showed a pronounced reduction in blood glucose, from ∼400 to ∼200 mg/dl, after a single oral gavage of D-allulose (Fig. 1H). Notably, the plasma insulin levels were not elevated but instead decreased significantly at 1 to 2 hours post-administration (Fig. 1I). The blood glucose-lowering effect of D-allulose was completely abolished by pretreatment with the GLP-1 receptor antagonist exendin(9–39) (Fig. 1J–1L). Moreover, intraperitoneal administration of D-allulose, which does not stimulate GLP-1 secretion (17), failed to increase portal GLP-1 levels and lower blood glucose (Fig. 1M–N).

**Figure 1.**
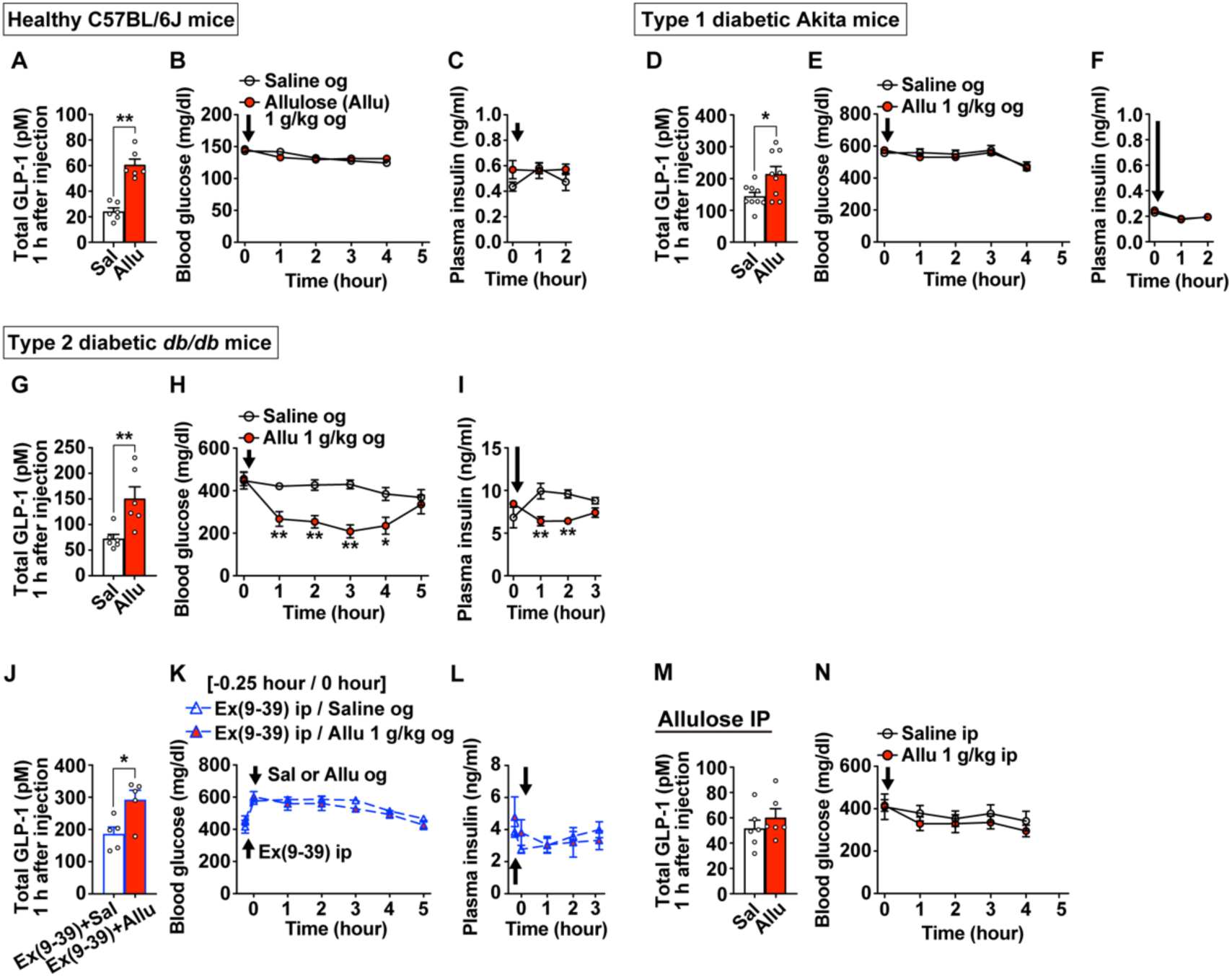
D-allulose-induced GLP-1 release enhances insulin action via GLP-1 receptor signaling in type 2 diabetic mice, but not in lean or type 1 diabetic mice. (**A–C**) Healthy C57BL/6J mice (n = 5–6) fasted for 4 hours received oral gavage (og) administration of saline or D-allulose at 1 g/kg. Plasma total GLP-1 levels in the portal vein were measured 1 hour after administration (**A**), along with sequential blood glucose levels (**B**) and plasma insulin levels (**C**). (**D–F**) Type 1 diabetic Akita mice (n = 5 or 9) fasted for 4 hours received oral D-allulose at 1 g/kg. Plasma total GLP-1 levels (**D**), sequential blood glucose levels (**E**), and plasma insulin levels (**F**) were assessed. (**G–I**) Type 2 diabetic *db/db* mice (n = 5–6) fasted for 4 hours received oral D-allulose at 1 g/kg. Plasma total GLP-1 levels (**G**), sequential blood glucose levels (**H**), and plasma insulin levels (**I**) were measured. (**J–L**) *db/db* mice (n = 5) were pretreated with GLP-1 receptor antagonist Ex(9–39) at 600 nmol/kg administered intraperitoneally 15 min before oral gavage of D-allulose at 1 g/kg. Plasma total GLP-1 levels (**J**), sequential blood glucose levels (**K**), and plasma insulin levels (**L**) were measured. (**M, N**) *db/db* mice (n = 5) fasted for 4 hours received intraperitoneal (ip) injection of D-allulose at 1 g/kg. Plasma total GLP-1 levels (**M**) and sequential blood glucose levels (**N**) were measured. Data are presented as mean ± SEM. *p < 0.05, **p < 0.01 by unpaired t-test (**A**, **D**, **G**, **J**) or two-way ANOVA with Bonferroni’s post hoc test (**H and I**). Arrows indicate the timing of reagent administration. og: oral gavage.

We further extended these findings to diet–induced obese (DIO) mice, another model of hyperglycemia and hyperinsulinemia. Oral, but not intraperitoneal, administration of D-allulose increased portal GLP-1 levels and lowered blood glucose without altering plasma insulin levels (Supplementary Fig. 1 and 3A–C). Consistently, D-allulose failed to lower blood glucose in GLP-1 receptor knockout mice with DIO (Supplementary Fig. 3F–3H). Collectively, these findings indicate that D-allulose lowers blood glucose in T2DM mice by enhancing insulin action through gut-derived GLP-1 and GLP-1 receptor activation, independently of insulin secretion.

### D-allulose-induced GLP-1 enhances insulin sensitivity once plasma insulin exceeds a certain threshold level

D-allulose-induced gut GLP-1 release lowered blood glucose only in hyperinsulinemic models, including *db/db* and DIO mice, but not under condition of low circulating insulin levels such as healthy and Akita mice (Fig. 1, Supplementary Fig. 2 and 3), suggesting that sufficient circulating insulin is required for this effect. To further test this possibility, circulating insulin levels in hypoinsulinemic Akita mice were elevated by intraperitoneal insulin injection. Insulin administration alone failed to significantly lower blood glucose (Fig. 2A), in consistent with the report of insulin resistance in this model (30). Notably, pretreatment with D-allulose recruited the action of exogenous insulin to significantly lower blood glucose (Fig. 2A–2B). This effect was completely abolished by pretreatment with the GLP-1 receptor antagonist (Fig. 2C–2D).

**Figure 2.**
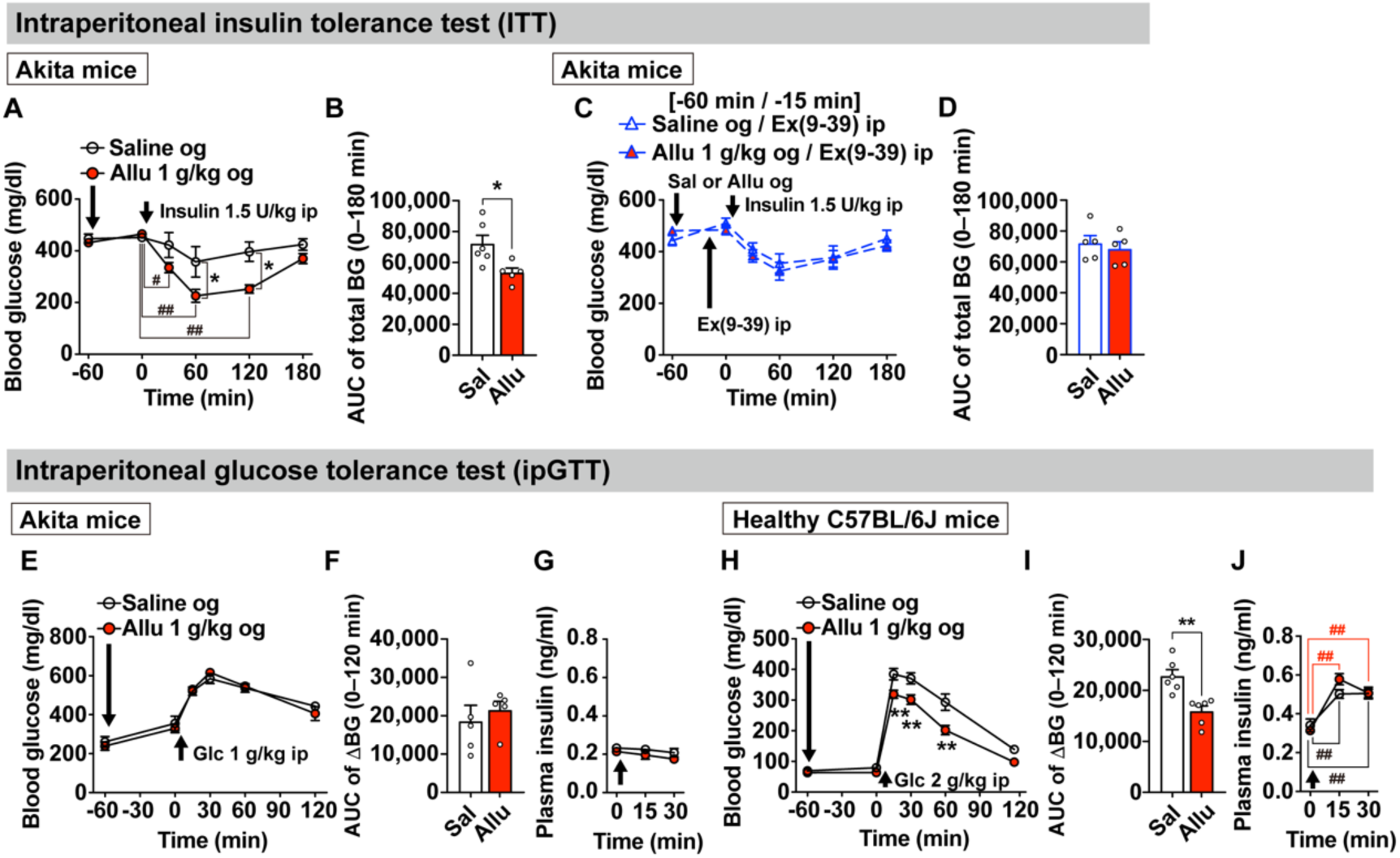
Suprathreshold levels of exogenous or endogenous insulin are required for blood glucose lowering by D-allulose induced GLP-1 release. (**A–D**) Insulin tolerance test in Akita mice fasted for 4 hours (n = 5–6). Mice received either oral gavage (og) of D-allulose at 1 g/kg or saline 1 hour before intraperitoneal (ip) injection of insulin at 1.5 U/kg. In panels **C** and **D**, GLP-1 receptor antagonist Ex(9–39) was administered intraperitoneally at 600 nmol/kg 15 min before insulin injection. Sequential blood glucose levels (**A, C**) and their area under the curve (AUC) from 0 to 180 min (**B, D**) were assessed. (**E–J**) Ip glucose tolerance test (ipGTT) in Akita mice and healthy C57BL/6J mice fasted for 16 hours (overnight). Akita mice (n = 5) received ip injection of glucose at 1 g/kg (**E–G**) and healthy C57BL/6J mice (n = 6) received 2 g/kg glucose (**H–J**). D-allulose at 1 g/kg or saline was administered orally 1 hour before glucose injection. Sequential blood glucose levels (**E**, **H**), AUC from 0 to 120 min (**F**, **I**), and plasma insulin levels (**G**, **J**) were measured. Data are presented as mean ± SEM. *p < 0.05, **p < 0.01 by unpaired t-test (**B, I**); two-way ANOVA with Bonferroni’s post hoc test vs. saline (**A, H**); #p < 0.05, ##p < 0.01 by two-way ANOVA with Tukey’s post hoc test vs. 0 min of each group (**A**); ##p < 0.01 by two-way ANOVA with Dunnett’s post hoc test vs. 0 min of each group (**J**). Arrows indicate timing of reagent administration. Glc: glucose. og: oral gavage.

During an intraperitoneal glucose tolerance test (ipGTT), oral gavage of D-allulose failed to improve glucose tolerance in Akita mice with impaired glucose-stimulated insulin secretion (GSIS) (Fig. 2E–2G). In contrast, in healthy mice exhibiting robust GSIS, pretreatment with D-allulose significantly improved glucose tolerance without affecting the magnitude of insulin secretion between groups following glucose challenge (Fig. 2H–2J). Together, these results demonstrate that D-allulose-induced GLP-1 enhances insulin sensitivity when circulating insulin levels exceed a critical threshold.

### D-allulose-induced GLP-1 cooperatively activates the common hepatic branch of vagal afferents with pancreatic insulin

Based on our previous finding that GLP-1 and insulin synergistically and additively activate a subpopulation of isolated nodose ganglion neurons responsive to both hormones (31), we examined whether D-allulose-induced GLP-1 cooperates with insulin to activate vagal sensory neurons *in vivo*, using pERK1/2 immunoreactivity as a marker of neuronal activation (17,32–34). Administration of either D-allulose (GLP-1 secretagogue) or glibenclamide (insulin secretagogue) alone approximately doubled the number of pERK1/2-positive neurons in both left and right nodose ganglia (NGs), and this activation was further augmented by their combined administration (Fig. 3A–3F). There was no appreciable difference in activation between the left and right NGs.

**Figure 3.**
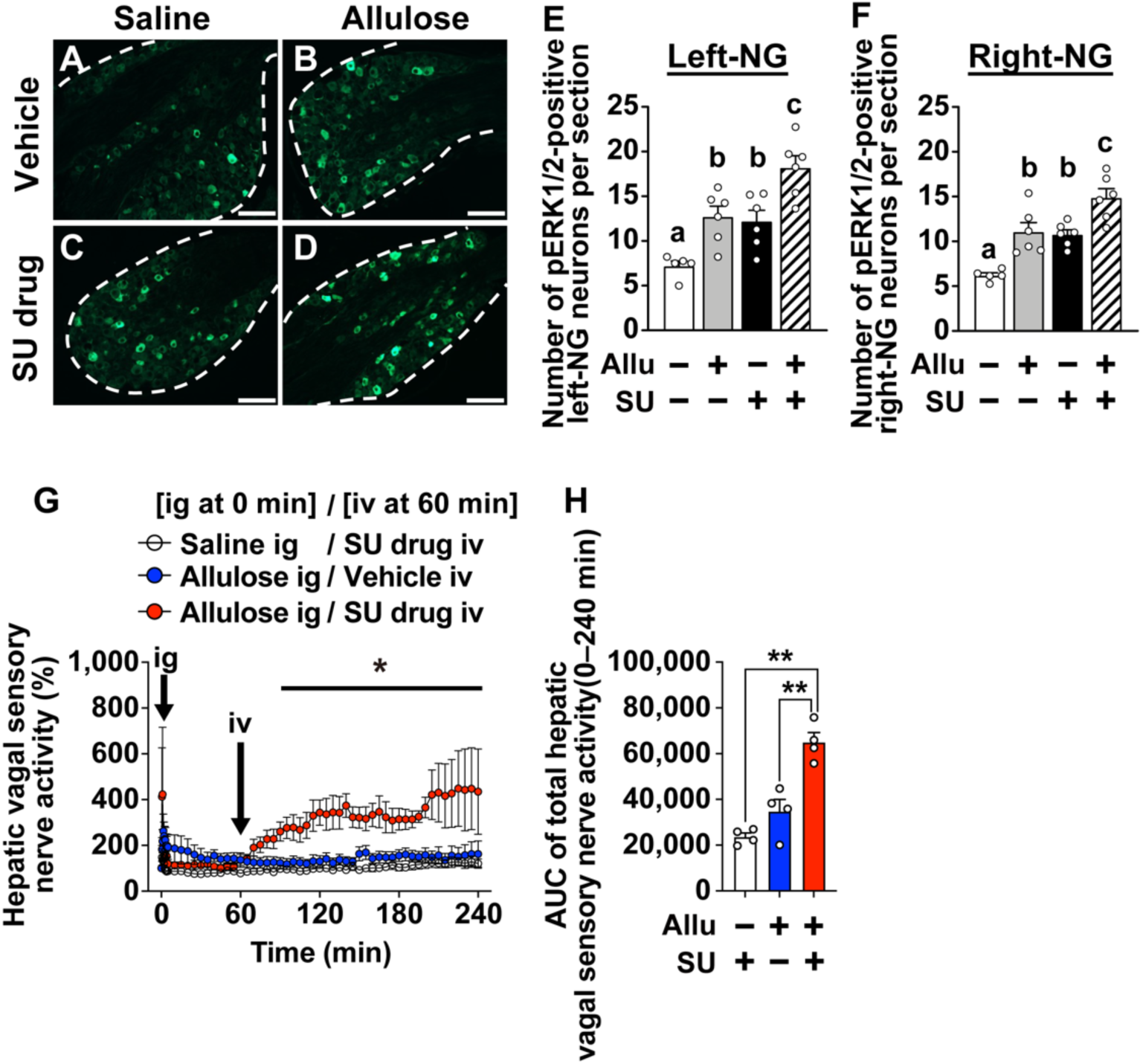
D-allulose-induced GLP-1 and pancreatic insulin cooperatively activates the common hepatic branch of vagal afferent nerves. (**A–E**) C57BL/6J mice received oral gavage (og) of D-allulose at 1 g/kg or saline at 30 min and ip injection of sulfonylurea drug glibenclamide (SU drug) at 1.5 mg/kg or vehicle at 15 min prior to transcardial perfusion. Phosphorylated ERK1/2 (pERK1/2) immunohistochemistry was performed on both left and right nodose ganglion (NG). Representative images of pERK1/2-positive neurons in left NG (green) are shown (**A–D**). Scale bars, 100 µm. Quantification of pERK1/2-positive neurons in the left (**E**) and right (**F**) NG. n = 5–6. Allu(–) and Allu(+) indicate oral gavage of saline or D-allulose, respectively; SU(–) and SU(+) indicate ip injection of vehicle or glibenclamide, respectively. (**G and H**) Neural activity of the common hepatic branch of vagal sensory nerve was recorded in anesthetized C57BL/6J mice (**G**). Mice received intragastric (ig) administration of D-allulose at 1 g/kg or saline at 0 min, followed by intravenous (iv) injection of glibenclamide at 1.5 mg/kg or vehicle at 60 min. Time-course changes in hepatic vagal sensory nerve activity (**G**) and its total AUC during 0 to 240 min (**H**). n = 4. Allu(–) and Allu(+) indicate ig injection of saline or D-allulose, respectively; SU(–) and SU(+) indicate iv injection of vehicle or glibenclamide, respectively. Data are expressed as means ± SEM. Different alphabet letters indicate p < 0.05 by one-way ANOVA followed by Tukey’s test in **E** and **F**, *p < 0.05 by two-way ANOVA followed by Dunnett’s test in **G** (vs. saline ig/glibenclamide iv group), and **p < 0.01 by one-way ANOVA followed by Tukey’s test in **H**. Arrows in the figures indicate the timing of administration of each reagent. og: oral gavage.

Next, we performed electrophysiological recordings of the common hepatic branch of the vagal afferents in anesthetized mice, a nerve critical for sensing nutrients and gastrointestinal-pancreatic hormones (15,35,36). Intragastric (ig) administration of D-allulose alone had minimal effects on activation of vagal sensory nerves (Fig. 3G, blue trace), whereas subsequent intravenous (iv) administration of glibenclamide 60 min after D-allulose resulted in a pronounced and sustained increase in firing frequency (Fig. 3G, red trace). Quantification of neuronal activity over 240 min revealed that combined D-allulose and glibenclamide administration elicited greater activation of the hepatic branch of vagal afferents compared to either treatment alone (Fig. 3H). These results indicate that vagal afferent nerves are strongly activated when both GLP-1 and insulin are simultaneously augmented.

### Cooperative action of gut–derived GLP-1 and pancreatic insulin on the common hepatic branch of vagal afferents enhances insulin-induced glucose lowering

In healthy mice, intraperitoneal administration of glibenclamide significantly lowered blood glucose, and this effect was markedly potentiated by pretreatment with D-allulose (Fig. 4A–B). Importantly, D-allulose pretreatment did not affect glibenclamide-induced insulin secretion (Fig. 4C). This potentiating effect of D-allulose was completely blunted in *Glp1r* knockout mice, despite preserved D-allulose–induced GLP-1 secretion and glibenclamide–induced insulin secretion (Fig. 4D–4G). To assess the involvement of vagal sensory nerves, we examined mice with chemical denervation of systemic peripheral sensory nerves using capsaicin (Fig. 4H–4K), or selective chemical denervation of the common hepatic branch of the vagus nerve (Fig. 4I–4O). In both models, D-allulose failed to enhance glibenclamide–induced glucose lowering, although D-allulose-induced GLP-1 release and glibenclamide–induced insulin secretion remained intact (Fig. 4H–4O).

**Figure 4.**
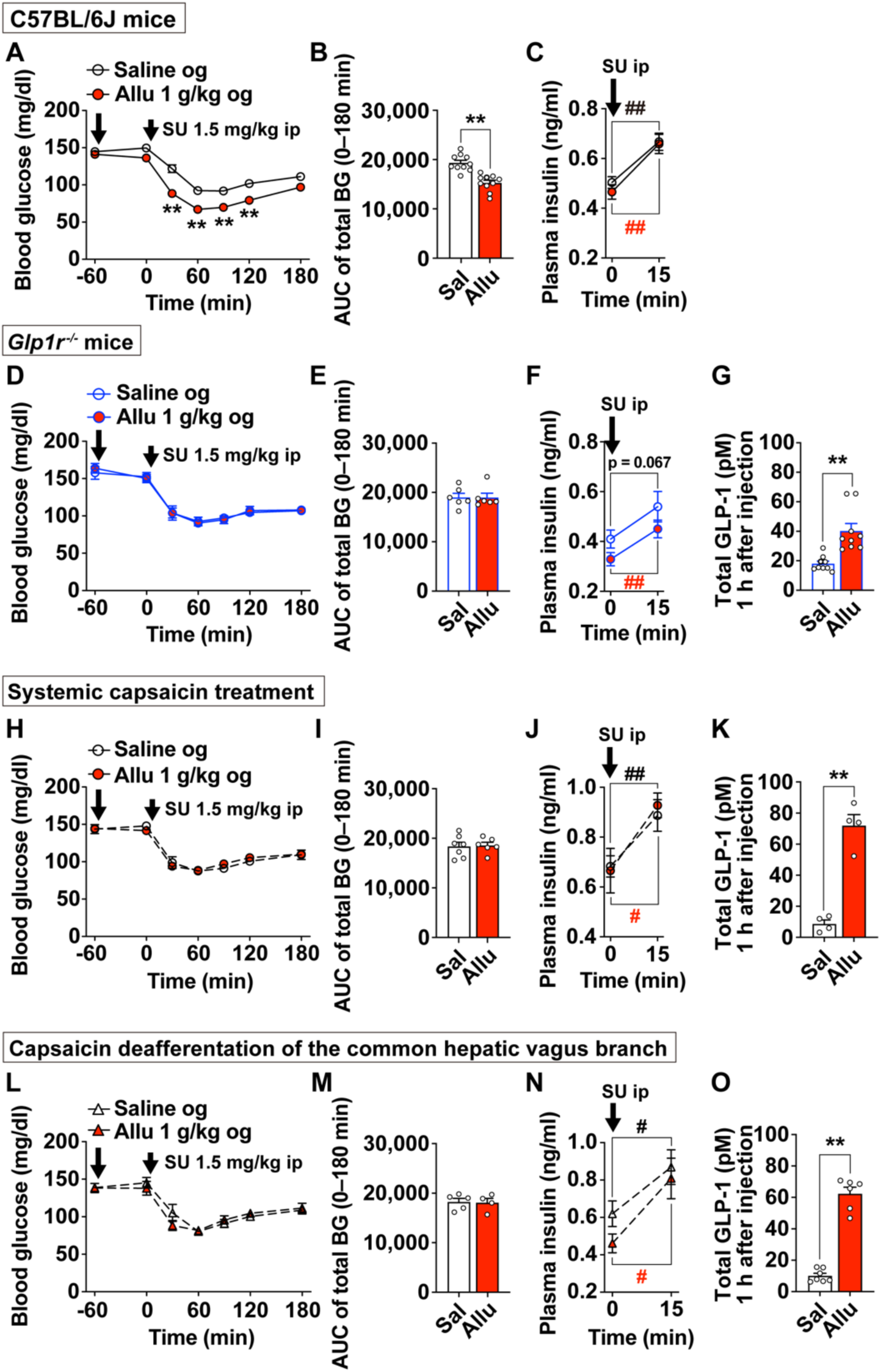
D-allulose enhances sulfonylurea-induced glucose lowering via GLP-1 receptors and the common hepatic branch of the vagus nerve. (**A–C**) C57BL/6J mice (n = 10) received oral gavage of D-allulose at 1 g/kg or saline 60 min prior to ip injection of glibenclamide at 1.5 mg/kg. Blood glucose levels (**A**), its AUC of total blood glucose during 0 to 180 min (**B**), and plasma insulin levels at 0 min (baseline) or 15 min after SU drug injection (**C**) were measured. (**D–O**) *Glp1r* knockout mice (**D–G**, n = 6–9), systemic capsaicin-treated mice (**H–K**, n = 4–7), and mice with selective capsaicin deafferentation of the common hepatic branch of the vagus nerve (**L–O**, n = 4–7) were treated as described above. Blood glucose (**D, H, L**), its AUC (**E, I, M**) and plasma insulin (**F, J, N**) were assessed. Total GLP-1 concentration in the portal vein plasma (**G, K, O**) were individually measured 1 h after oral gavage of saline or D-allulose at 1 g/kg. Data are expressed as means ± SEM. **p < 0.01, *p < 0.05 by two-way ANOVA followed by Bonferroni’s test vs. saline group (**A**), unpaired *t*-test (**B**, **G**, **K**, **O**). ##p < 0.01, #p < 0.05 by paired *t*-test (**C**, **F**, **J**, **N**). Different letters indicate p < 0.05 by one-way ANOVA followed by Tukey’s test (**T** and **U**). Arrows in the figures indicate the timing of administration of each reagent. og: oral gavage.

Furthermore, D-allulose improved glucose tolerance in ipGTT and potentiated insulin–induced glucose lowering in insulin tolerance tests (ITT) in sham-operated mice (Supplementary Fig. 4A–4D). These effects were completely abolished by either surgical vagotomy (Supplementary Fig. 4E–4H) or chemical deafferentation (Supplementary Fig. 4I–4L) of the common hepatic branch of the vagus nerve, originating from the left NG. In contrast, transection of the subdiaphragmatic dorsal trunk, which arises from the right NG, did not affect the action of D-allulose in ipGTT or ITT (Supplementary Fig. 4M–4P).

Together, these results demonstrate that gut GLP-1 and pancreatic insulin cooperatively enhance systemic insulin sensitivity via the left–sided common hepatic branch of vagal sensory nerves, thereby driving glucose lowering independently of further insulin secretion.

### GLP-1 receptor–expressing left vagal sensory neurons are essential for D-allulose–induced enhancement of insulin sensitivity

To determine whether GLP-1R–expressing vagal sensory neurons mediate the cooperative effects of GLP-1 and insulin, we selectively expressed the inhibitory DREADD hM4Di in GLP-1R–expressing neurons by microinjecting AAV2-hSyn-DIO-hM4D(Gi)-mCherry into either the left or right NG of *Glp1r-ires-Cre* mice (Fig. 5A–5C, 5L–5N). Under vehicle–treated conditions, D-allulose potentiated glibenclamide-induced glucose lowering and improved glucose tolerance during ipGTT in *Glp1r-Cre* mice with left NG–specific hM4Di expression (Fig. 5D, 5E, 5H, 5I). This effect was completely abolished by pretreatment with the hM4Di ligand clozapine *N*-oxide (CNO) (Fig. 5F, 5G, 5J, 5K). In contrast, mice with hM4Di expression in the right NG retained D-allulose–induced enhancement of glucose lowering and glucose tolerance, regardless of CNO administration (Fig. 5O–5V). These results indicate that left-sided GLP-1R–expressing vagal afferents are critical for the cooperative enhancement of insulin sensitivity by GLP-1 and insulin.

**Figure 5.**
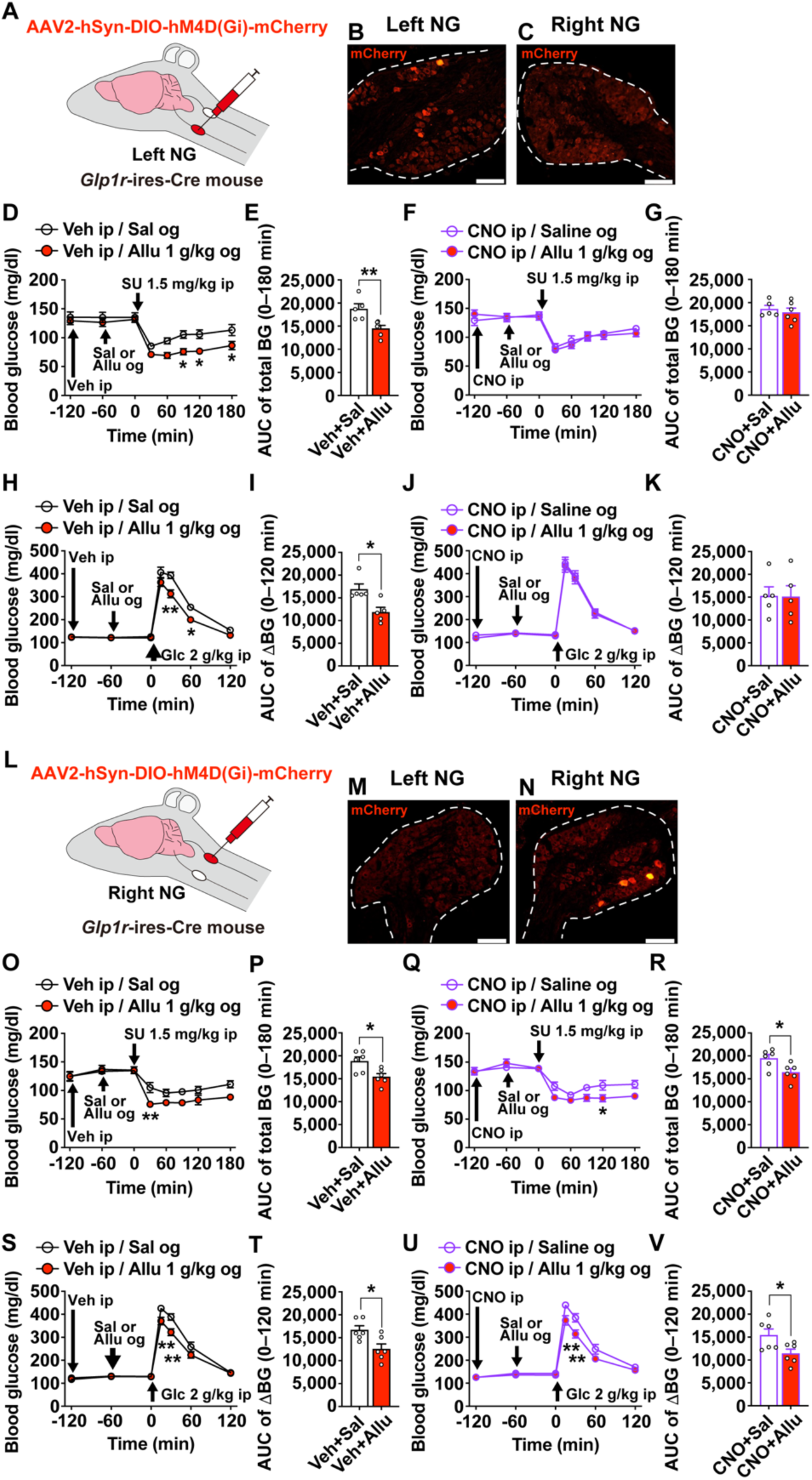
Left-sided, but not right-sided, GLP-1R-expressing vagal sensory nerves mediate the enhancement of insulin action by D-allulose-induced GLP-1 release. (**A**) AAV2-hSyn-DIO-hM4D(Gi)-mCherry was microinjected into the left nodose ganglion (NG) of *Glp1r-ires-Cre* mice. (**B** and **C**) Representative mCherry fluorescence in the left (**B**) and right (**C**) NG. (**D–K**) *Glp1r-Cre*^left^ ^NG-*hM4Di*^ mice were pretreated with clozapine-*N*-oxide (CNO) at 3 mg/kg (**F, G, J, K**) or vehicle (**D, E, H, I**) at −120 min relative to the start of the tolerance test. At −60 min, D-allulose 1 g/kg or saline was given orally. Finally, glibenclamide at 1.5 mg/kg (**D–G**) or glucose at 2 g/kg (**H–K**) was administered intraperitoneally at 0 min. Blood glucose levels (**D**, **F, H, J**) and its AUC during 0–180 min (**E**, **G, I, K**) were measured. (**L**) AAV2-hSyn-DIO-hM4D(Gi)-mCherry was microinjected into the right NG of *Glp1r-ires-Cre* mice. (**M** and **N**) Representative mCherry fluorescence in the left (**M**) and right (**N**) NG. (**O–V**) Same experimental procedure as in (**D–K**) was applied to *Glp1r-Cre^r^*^ight^ ^NG-*hM4Di*^ mice. Data are expressed as means ± SEM. **p < 0.01, *p < 0.05 by two-way ANOVA followed by Bonferroni’s test vs. saline group (**D, H, O, Q, S, U**), unpaired *t*-test (**E**, **I**, **P**, **R, T, V**). Arrows in the figures indicate the timing of administration of each reagent. og: oral gavage.

### IRS2-mediated insulin signaling in vagal sensory neurons enables the cooperative glucose lowering by GLP-1 and insulin

Insulin receptor substrate 2 (IRS2) is essential for insulin sensing in vagal sensory neurons (37). To examine the role of IRS2 in the cooperative action of GLP-1 and insulin on vagal sensory neurons, we generated *Phox2b-Cre:Irs2^flox/flox^*mice by crossing *Phox2b-Cre* mice (24) with *Irs2^flox/flox^* mice (22). *Irs2* mRNA expression was significantly reduced in both the NG and the nucleus tractus solitarius (NTS) in *Phox2b-Cre:Irs2^flox/flox^* mice, regions in which Cre recombinase is predominantly expressed (Fig. 6A). In contrast, *Irs2* mRNA expression remained unchanged in the hypothalamus, liver, skeletal muscle, and interscapular brown adipose tissue (Fig. 6A). Expression levels of other hormone receptors and insulin signaling molecules in NG were not altered (Supplementary Fig. 5).

**Figure 6.**
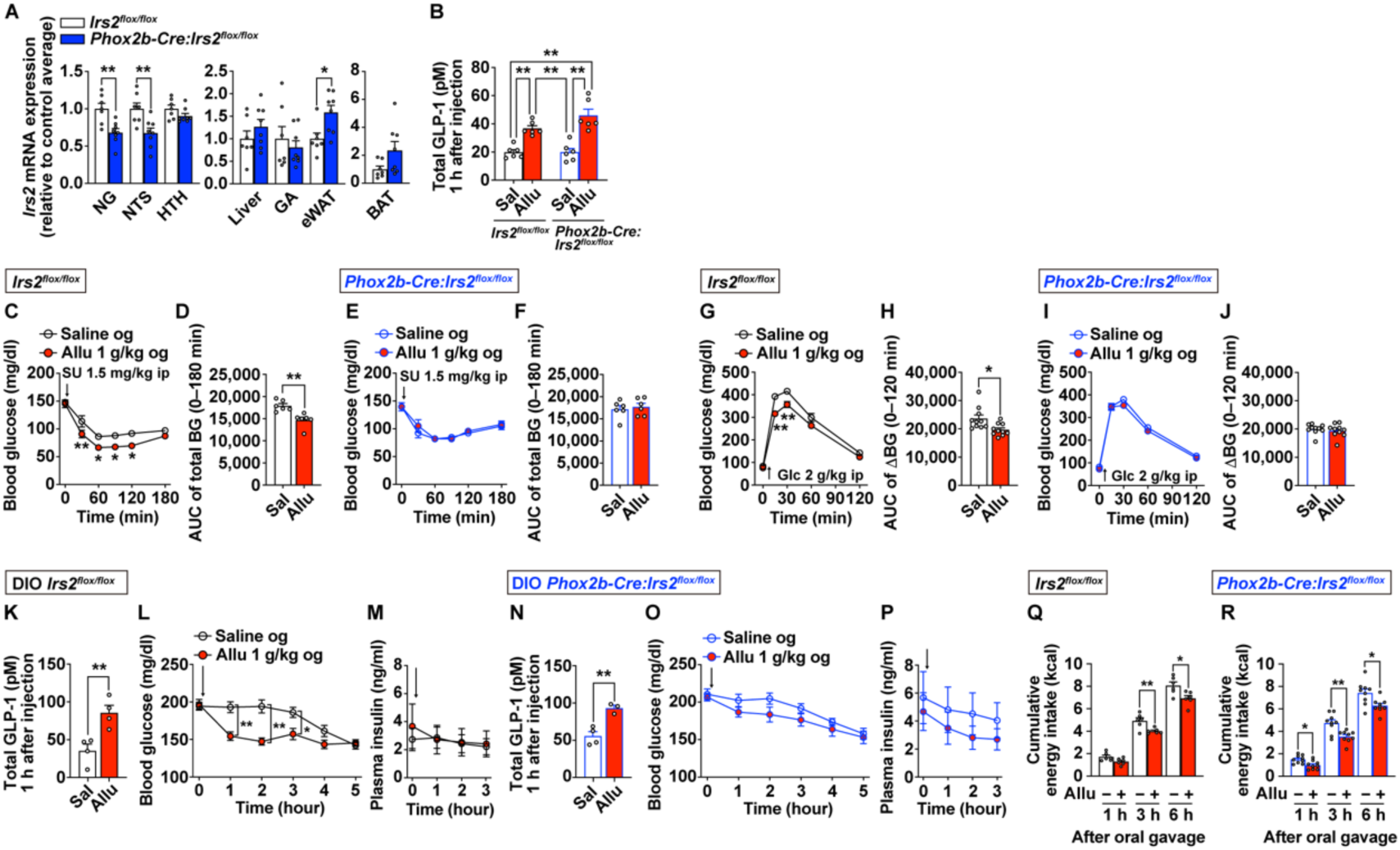
Vagal IRS2 signaling is essential for cooperative glucose lowering by GLP-1 and insulin, but not for their anorexigenic effects. (**A**) Vagal sensory nerve-preferential *Irs2* knockdown mice were generated by crossing *Phox2b-Cre* mice with *Irs2^flox/flox^* mice. Relative *Irs2* mRNA expression in several organs in lean *Irs2^flox/flox^*and lean *Phox2b-Cre:Irs2^flox/flox^* mice fed standard chow (n = 7–8). (**B**) Plasma GLP-1 levels in the portal vein 1 hour after oral gavage of D-allulose at 1 g/kg or saline in lean *Irs2^flox/flox^* and lean *Phox2b-Cre:Irs2^flox/flox^* mice (n = 6). (**C–F**) Blood glucose and its AUC during 0–180 min after ip injection of glibenclamide at 1.5 mg/kg in lean *Irs2^flox/flox^*(**C, D**) and lean *Phox2b-Cre:Irs2^flox/flox^* mice (**E, F**) fasted for 4 hours and pretreated with oral gavage of saline or D-allulose at 1 g /kg 1 h prior to glibenclamide injection (n = 9–10). (**G–J**) Blood glucose and its AUC during 0–120 min after ip injection of glucose at 2 g/kg in lean *Irs2^flox/flox^*(**G, H**) and lean *Phox2b-Cre:Irs2^flox/flox^* mice (**I, J**) fasted for 4 hours and pretreated with oral gavage of saline or D-allulose at 1 g/kg 1 h prior to glucose injection (n = 10). (**K-P**) Effect of oral gavage of saline or D-allulose at 1 g/kg on hyperglycemia in DIO *Irs2^flox/flox^* (**L**, **M**, n = 8) and DIO *Phox2b-Cre:Irs2^flox/flox^* mice (**O**, **P**, n = 7) fed a high-fat diet at least 2 months and fasted for 4 hours. Total GLP-1 concentration in the portal vein plasma were individually measured 1 h after oral gavage of saline or D-allulose at 1 g/kg (**K**, **N**, n = 3–4). (**Q, R**) Effects of oral gavage of saline or D-allulose at 1 g/kg on dark-phase feeding behavior in lean *Irs2^flox/flox^* and lean *Phox2b-Cre:Irs2^flox/flox^* mice fasted for 3 h (n = 6–9). Data are expressed as means ± SEM. **p < 0.01, *p < 0.05 by unpaired *t*-test (**A, D, H, K, O, Q, R**), one-way ANOVA followed by Tukey’s test (**B**) or two-way ANOVA followed by Bonferroni’s test vs. saline group (**C, G, L**). Arrows in the figures indicate the timing of administration of each reagent. og: oral gavage.

A single oral gavage of D-allulose similarly increased portal vein GLP-1 levels in both control *Irs2^flox/flox^* and *Phox2b-Cre:Irs2^flox/flox^*mice (Fig. 6B). In control mice, pretreatment of D-allulose enhanced glibenclamide–induced glucose lowering and improved glucose tolerance during ipGTT (Fig. 6C–6D, 6G–6H). However, both beneficial effects were completely blunted in *Phox2b-Cre:Irs2^flox/flox^*mice (Fig. 6E–6F, 6I–6J).

To further assess this mechanism under hyperinsulinemic conditions, control and *Phox2b-Cre:Irs2^flox/flox^* mice were fed a high-fat diet to generate DIO model. Oral gavage of D-allulose similarly increased portal GLP-1 levels in both DIO control and DIO *Phox2b-Cre:Irs2^flox/flox^* mice (Fig. 6K and 6N). D-allulose significantly lowered blood glucose levels without altering plasma insulin levels in DIO control mice (Fig. 6L–6M), whereas this effect was blunted in DIO *Phox2b-Cre:Irs2^flox/flox^* mice (Fig. 6O–6P). These results indicate that the elevated insulin by SU or glucose stimulation in lean mice or by obesity in DIO mice cooperates with the gut GLP-1 release by D-allulose to lower blood glucose levels, and that this cooperation requires IRS2 signaling in vagal sensory neurons.

We previously reported that D-allulose–induced GLP-1 release suppresses food intake via GLP-1 receptor–expressing vagal sensory neurons (17). However, in the present study, oral gavage of D-allulose suppressed food intake in both *Irs2^flox/flox^* and *Phox2b-Cre:Irs2^flox/flox^* mice (Fig. 6Q–6R). These findings show that IRS2 signaling in vagal sensory neurons is not required for the anorexigenic effect of D-allulose.

### GLP-1 release by D-allulose improves severe insulin resistance more effectively and through a mechanism distinct from pharmacological GLP-1R agonists

Finally, we compared the mechanisms of glucose regulation by gut–derived GLP-1 and pharmacological GLP-1 receptor agonists exendin-4 (Ex4). Ex4 was selected for this study because, among GLP-1 receptor agonists, it has the shortest half-life and exhibits short-term feeding suppression similar to that of endogenous GLP-1 (38,39).

Ex4 was administered at a minimal effective dose (0.5 nmol/kg) that robustly suppressed food intake within 30 min. Intraperitoneal Ex4 significantly increased plasma insulin levels and lowered blood glucose in both healthy and DIO mice (Supplementary Fig. 6A–6F), effects that were completely abolished in *Glp1r* knockout mice (Supplementary Fig. 6G–6L). Notably, chemical ablation of peripheral sensory nerves using capsaicin did not affect Ex4–induced insulin secretion and glucose lowering (Supplementary Fig. 6M–6O). These results clearly demonstrate that the mechanism of glucose regulation by GLP-1R agonists is fundamentally distinct from that of gut–derived GLP-1 induced by D-allulose.

Compared with DIO mice (Supplementary Fig. 3), *db/db* mice exhibited more severe hyperglycemia and hyperinsulinemia (Fig. 7A–7D), indicating a higher degree of insulin resistance. In *db/db* mice, a single oral gavage of D-allulose markedly and rapidly improved severe hyperglycemia, together with a tendency to reduce plasma insulin levels (Fig. 1H, 1I, 7A and 7B). Conversely, Ex4 robustly increased insulin secretion but failed to significantly lower blood glucose, possibly due to the severe insulin resistance (Fig. 7C and 7D) (3). These results demonstrate that, in a single-dose setting under conditions of severe insulin resistance, stimulation of endogenous GLP-1 release by D-allulose achieves superior glycemic control compared with pharmacological GLP-1 receptor activation by Ex4.

**Figure 7.**
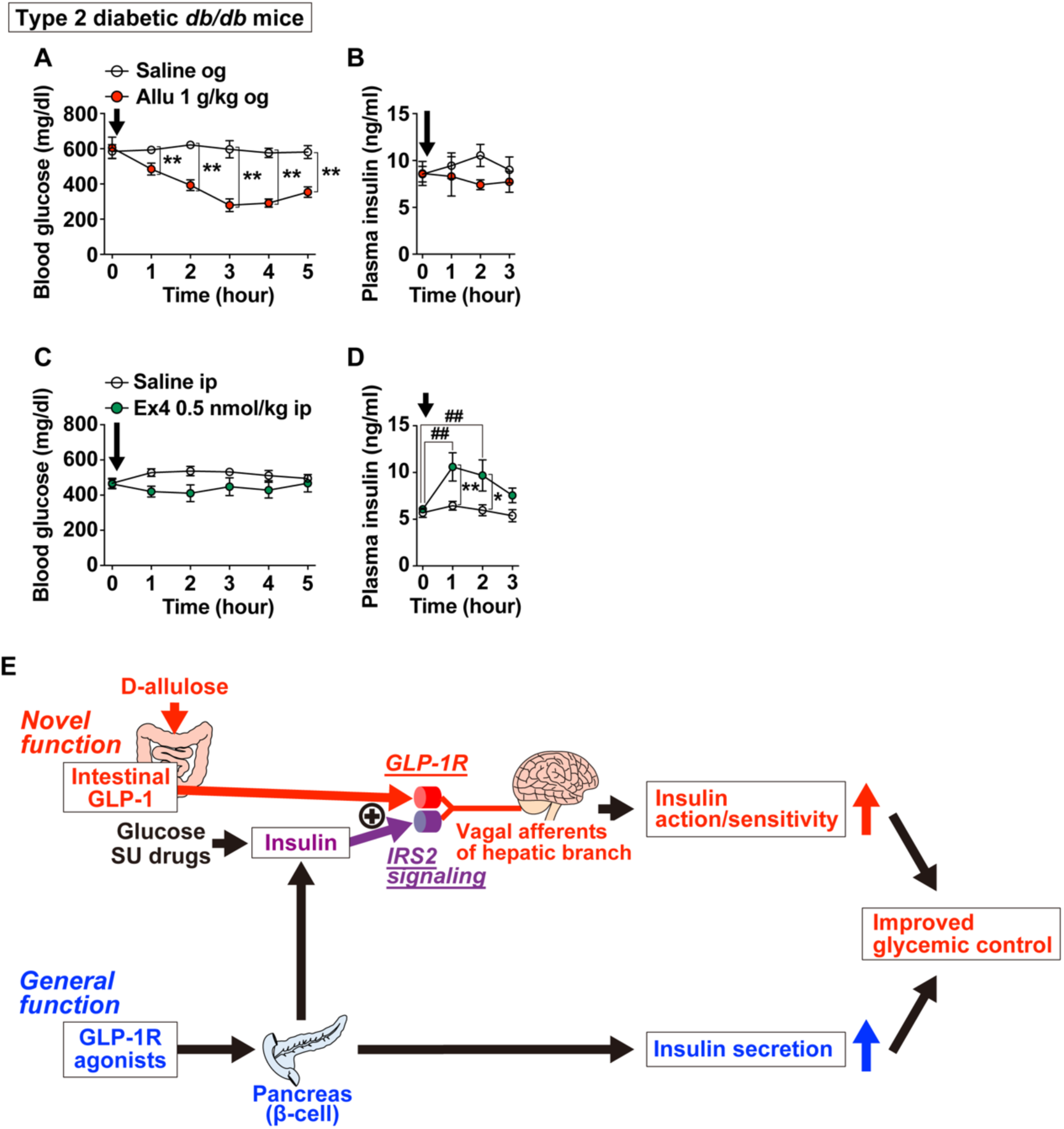
Comparison of hyperglycemia-improving effects of GLP-1–stimulating D-allulose vs. GLP-1 receptor agonists in type 2 diabetes and proposed mechanism of action. (**A–D**) Sequential blood glucose and plasma insulin levels after single administration of D-allulose (1 g/kg, og, n = 5-6, **A and B**) or exendin-4 (Ex4, 0.5 nmol/kg, ip, n = 5-6, **C and D**) in T2DM *db/db* mice fasted for 4 hours. Arrows in the figures indicate the injection timing of D-allulose or Ex4. Data are expressed as means ± SEM. **p < 0.01, *p < 0.05 by two-way ANOVA followed by Bonferroni’s test vs. saline group. ##p < 0.01 by two-way ANOVA followed by Dunnett’s test vs. 0 h of each group. og: oral gavage. (**E**) Graphical summary of this study. Proposed mechanism for the action of intestinal GLP-1 release induced by D-allulose, representing a novel route of GLP-1 function identified in this study. Cooperative action of GLP-1 release and insulin release on the common hepatic branch of vagal afferents co-expressing GLP-1R and IRS2 is linked to enhanced insulin action/sensitivity. This mechanism is driven when endogenous insulin levels are elevated, such as during glucose ingestion, sulfonylurea administration, or under basal hyperinsulinemic conditions in type 2 diabetes, leading to suppression of postprandial glucose excursions and improvement of hyperglycemia. In contrast, pharmacological levels of GLP-1 or GLP-1 receptor agonists lower blood glucose by stimulating insulin secretion through activation of GLP-1Rs expressed on pancreatic β-cells (general function of GLP-1).

## DISUCUSSION

In the present study, we identified a novel glucose–lowering mechanism whereby D-allulose–induced gut–derived GLP-1 enhances insulin sensitivity to improve hyperglycemia through GLP-1 receptors– and IRS2–expressing left–sided vagal sensory neurons (Fig. 7E). Using non-calorie GLP-1 secretagogue D-allulose, we demonstrated that stimulation of gut GLP-1 release alone does not lower blood glucose under conditions of low or normal circulating insulin. In contrast, when insulin levels were elevated by sulfonylureas, glucose administration, or under hyperinsulinemic diabetic states, gut–derived GLP-1 release lowered blood glucose by enhancing insulin sensitivity without further increasing insulin secretion. This glucose–lowering effect required cooperative activation of vagal sensory neurons by both GLP-1 and insulin, and its effect was completely abolished by either chemogenetic inhibition of GLP-1R–expressing vagal sensory neurons or genetic deletion of IRS2 in these neurons. Furthermore, we found that this glucose–regulatory pathway is mediated specifically by the left–sided common hepatic branch of the vagal sensory nerves, revealing a lateralized neural control of glucose metabolism. Notably, in T2DM mice with severe insulin resistance, D-allulose–induced gut GLP-1 secretion exerted superior glucose–lowering effects compared to the GLP-1 receptor agonist Ex4. Together, these findings reveal a previously unrecognized physiological role of intestinal GLP-1 in improving systemic insulin sensitivity via a vagal afferent pathway, which is mechanistically distinct from pharmacological GLP-1 receptor agonism (Fig. 7E).

The common hepatic branch of the vagus nerve plays a critical role in sensing postprandial signals, and has been recognized as a major neural target of GLP-1 (15,35,36,40). Previous studies have shown that GLP-1 acts alone on vagal sensory neurons to enhance insulin secretion from pancreatic β-cells, known as the "neuroincretin" effect (14,15,35). In contrast, our findings demonstrate that gut–derived GLP-1 activates vagal sensory neurons more robustly when acting in concert with insulin than when acting alone, leading not to further insulin secretion but to enhanced systemic insulin sensitivity. Our previously study, using cytosolic calcium imaging of isolated single NG neurons demonstrated that GLP-1 or insulin at locally high concentrations (10^-8^–10^-7^ M) activates a subclass of these neurons via GLP-1R or IRS2, respectively (31,37), and that the neuronal populations responsive to GLP-1 and insulin largely overlap (31). At physiological concentrations (10^-9^ M), GLP-1 or insulin alone only weakly activated a small subclass of these neurons, whereas their co-application enhanced the Ca^2+^ response and recruited neurons unresponsive to either hormone alone (31,41). Consistent with this, the current *in vivo* data show that D-allulose–induced GLP-1 secretion and sulfonylurea-induced insulin release together activated vagal afferents in the common hepatic branch more robustly than either stimulus alone (Fig. 3G and 3H). These results indicate the existence of a distinct neuronal subclass that is activated exclusively by the combined presence of GLP-1 and insulin (31,41). Furthermore, our results reveal that IRS2 signaling in vagal sensory neurons is required for enhancement of insulin sensitivity, but not for the anorexigenic action of gut–derived GLP-1 (Fig. 6C–6P). The present physiological findings suggest that distinct neuronal subclasses exist, one activated by GLP-1 alone and another activated by the cooperative action of GLP-1 and insulin, with each group potentially controlling different functions such as feeding behavior, insulin secretion, or insulin sensitivity.

Vagal sensory neurons are pseudounipolar neurons whose cell bodies reside in the left and right nodose ganglia. These neurons project to their respective ipsilateral NTS in the brainstem, whereas their peripheral innervation patterns differ due to the asymmetric arrangement of visceral organs (40). Recent studies have revealed that this left–right asymmetry extends beyond anatomy and includes functional specializations (17,27,42–44). In this study, we found that the enhancement of systemic insulin sensitivity by gut–derived GLP-1 and insulin required afferent input from the left nodose ganglion–derived common hepatic branch. In contrast, D-allulose–induced intestinal GLP-1 secretion activated not only left–sided but also right–sided vagal afferents (Fig. 3A–3F). The physiological role of the right-sided vagal activation by gut–derived GLP-1 remains unclear and require future investigation.

The pathogenesis of T2DM involves two major factors, impaired insulin secretion and reduced insulin action. Pharmacological strategies to enhance insulin secretion have advanced considerably and have been continuously refined in terms of efficacy and safety. However, therapeutic approaches targeting insulin resistance remain limited. We have previously demonstrated that gut–derived GLP-1 release enhances systemic insulin sensitivity and glucose tolerance via activation of vagal sensory neurons (17,18,25). In the present study, we identify a previously unrecognized mechanism by which gut–derived GLP-1, acting in concert with insulin, enhances systemic insulin sensitivity through GLP-1R– and IRS2–expressing left–sided vagal sensory neurons. This newly identified action of gut–derived GLP-1 may help preserve β-cells by reducing the need for excessive postprandial insulin secretion, highlighting its importance for long-term metabolic control and its therapeutic potential.

Indeed, earlier clinical studies have demonstrated that augmenting endogenous GLP-1 secretion prior to carbohydrate intake by altering the meal sequence can suppress postprandial glucose excursions without increasing plasma insulin levels (45). The authors originally attributed the blunted postprandial rise in plasma insulin levels to the smaller postprandial increase in blood glucose. However, based on our findings, it is plausible that gut GLP-1 release enhances insulin sensitivity via vagal pathway, thereby contributing to the observed glycemic benefit. Additionally, recent studies have demonstrated that D-allulose stimulates GLP-1 secretion in humans (46), and that the addition of a fixed amount of D-allulose to meals significantly reduces postprandial glucose excursions (47). Importantly, D-allulose–induced GLP-1 secretion did not cause hypoglycemia (Fig. 1B), highlighting a clinically relevant safety feature. These human findings raise the possibility that the glycemic effects of D-allulose may also involve the mechanism demonstrated in the present study. Further investigation of dietary strategies, particularly the timing of D-allulose intake as a GLP-1 releaser relative to meal initiation, may enable improved postprandial glycemic control and support the development of therapeutic approaches, including drug development, nutritional guidance, and functional diet design, for individuals with or at risk for T2DM.

One of the most striking findings of this study is that a single administration of D-allulose improved hyperglycemia in T2DM *db/db* mice more rapidly and effectively than a GLP-1 receptor agonists (Fig. 7A–7D). While GLP-1 receptor agonists primarily lower blood glucose by stimulating insulin secretion from pancreatic β-cells (11,12), D-allulose–induced GLP-1 release enhances insulin sensitivity via the vagal pathway, representing fundamentally approach to T2DM treatment. Although the precise mechanisms remain unclear, several studies have reported that GLP-1 receptor agonists, including exendin-4, do not elicit significant activation of vagal sensory neurons (Supplementary Fig. 6M–6O) (23,48). In contrast, accumulating evidence, including the present study, demonstrates that endogenously released GLP-1 robustly activates vagal afferent pathways, thereby engaging gut–brain neural circuits that regulate systemic insulin sensitivity. In addition, a diverse array of naturally occurring dietary components, including nutrients and low- or non-caloric food ingredients such as D-allulose and polyphenols, have been identified as potent inducers of GLP-1 secretion (17,49–51). Therefore, integrating pharmacological therapy with dietary strategies that boost intestinal GLP-1 release and enhance insulin sensitivity may lead to more effective and sustainable approach to glycemic control in T2DM. Moreover, given the growing concern over glycemic rebound following discontinuation of GLP-1 receptor agonists (52), targeting the gut–vagal–brain axis may provide a novel adjunctive or alternative strategy to maintain long-term metabolic benefits.

## ACKNOWLEDGMENTS

The authors thank Dr. Hiroshi Inoue (Kanazawa University) for kindly providing live Phox2b-Cre mice (Jackson Laboratory, strain no. 016223), Dr. Yasuhiko Minokoshi (Sugiyama Jogakuen University) for valuable scientific input on glucose metabolism, and Dr. Ken-ichiro Nakajima (Nagoya University) and Dr. Kunio Kondoh (Tottori University) for advice on DREADD experiments using AAV vectors. We also thank Dr. Yuta Masuda, Rika Kitano, Tenko Shimizu (Kyoto Prefectural University) for technical assistance with DREADD experiments, as well as all laboratory members including Yudai Sugiyama and Jun Yoshitake (Kyoto Prefectural University) for their experimental support.

## AUTHOR CONTRIBUTIONS

K.O., T.Y. and Y.I. developed concept and designed the study. K.O., M.T., H.I., W.O. and Y.I. performed experiments and analyzed data. K.O. C.A., and Y.I. contributed to establishing the microinjection method into the nodose ganglion in mice. N.K. and D.J.D. developed and provided the key genetically modified mouse strain. K.O., M.T., T.Y., and Y.I. prepared figures, interpreted the results of the experiments, and drafted the manuscript. All of the authors edited the final draft. All authors have read and agreed to the published version of the manuscript.

## SUPPLEMENTARY METHODS

### Induction of insulin-deficient type 1 diabetes in mice using streptoxotocin (STZ)

STZ was administered to ablate pancreatic β-cells, leading to impaired insulin secretion. Mice received intraperitoneal injections of STZ at a total dose of 400 mg/kg, administered as two doses of 200 mg/kg on Day 1 and Day 3. On Day15, plasma insulin levels were measured using an insulin ELISA kit (MS303; Morinaga, Yokohama, Japan), confirming that the mice had insulin levels below the detection limit of the kit and were subsequently used for behavioral experiments.

### Measurement of glucagon in portal vein plasma

Plasma glucagon concentrations were measured using a commercially available ELISA kit (Mercodia AB, Uppsala). Blood collection and plasma preparation were performed as described for the "Measurement of GLP-1 in portal vein plasma".

### Quantitative PCR analysis

Primers sequences were as follows:

Irs1-Forward: 5’-TGGACATCACAGCAGAATGAAGA-3’
Irs1-Reverse: 5’-GACGTGAGGTCCTGGTTGTG-3’
Irs2-Forward: 5’-CACAGTCGTGAAAGAGTGAAGC-3’
Irs2-Reverse: 5’-GTTGGTCGGAAACATGCCAA-3’
Glp1r-Forward: 5’-TGTTGGCTTCAGACACTTGC-3’
Glp1r-Reverse: 5’-TACATCCACTTGAGGGCAGC-3’
Insr-Forward: 5’-ATGGGCTTCGGGAGAGGAT-3’
Insr-Reverse: 5’-GGATGTCCATACCAGGGCAC-3’
Cckar-Forward: 5’-AAAAGAATGGCAGTCTGCAGTG-3’
Cckar-Reverse: 5’-TGATAACCAGCGTGTTCCCC-3’

## SUPPLEMENTARY FIGURES AND FIGURE LEGENDS

**Supplementary Fig. 1.**
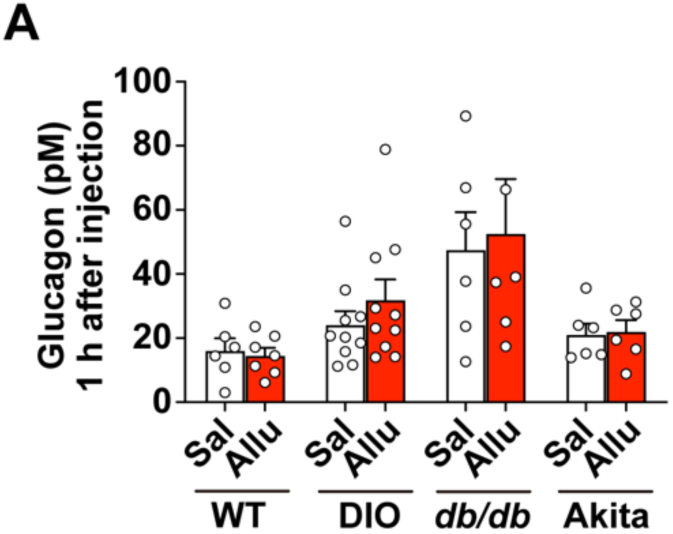
Oral gavage of D-allulose dose not alter plasma glucagon levels in several model mice. (**A**) Healthy C57BL/6J (WT, n = 6–7), diet-induced obese (DIO, n = 10), *db/db* (n = 5–6), and Akita mice (n = 6) fasted for 4 hours received oral gavage of D-allulose at 1 g/kg. Plasma glucagon levels in the portal vein remained unchanged across all groups. Data are expressed as means ± SEM. Not significant by unpaired *t*-test.

**Supplementary Fig. 2.**
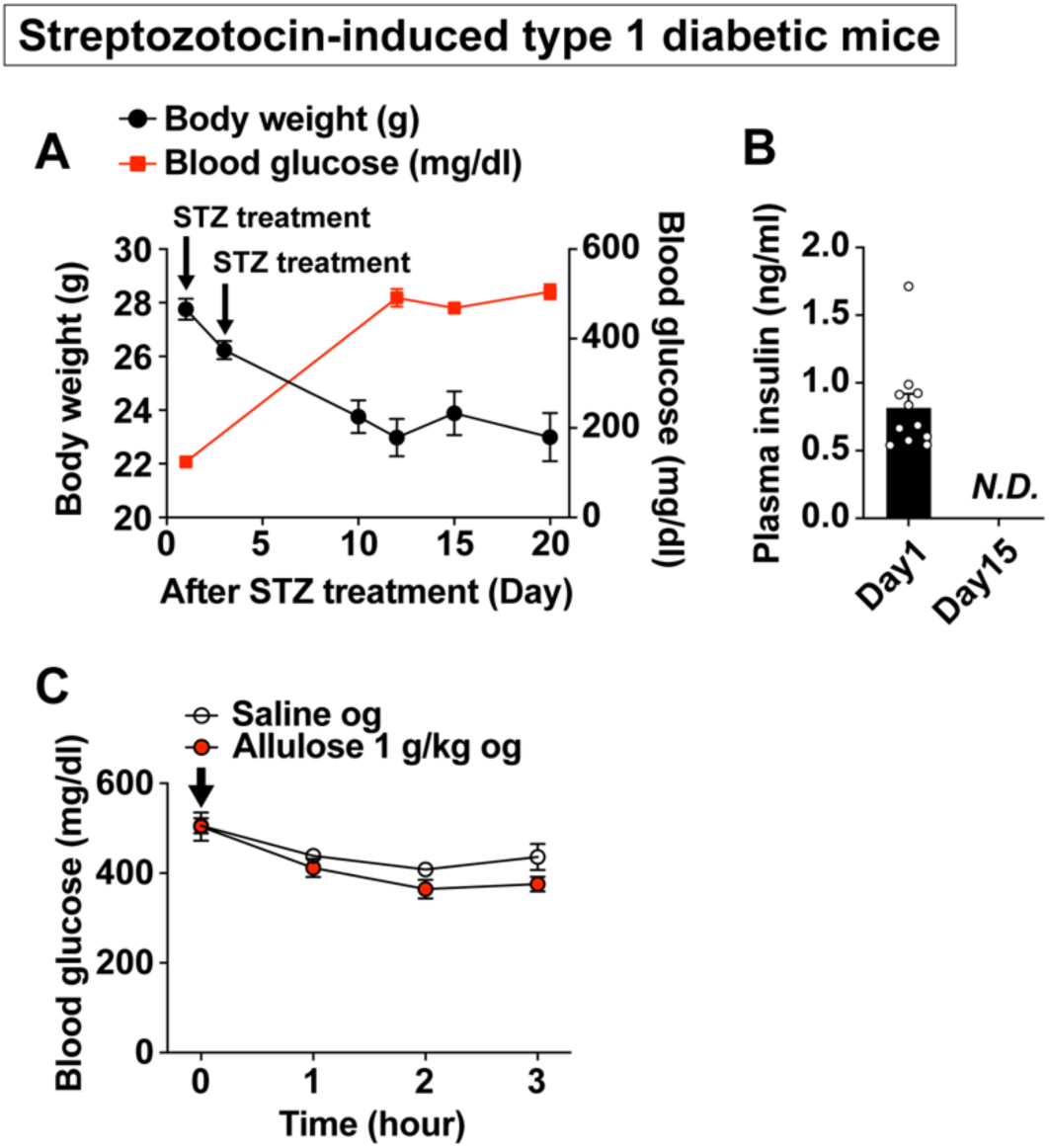
D-allulose fails to improve hyperglycemia in streptozotocin (STZ)-induced type 1 diabetic mice with complete β-cell ablation. (**A**) Time course of body weight (black line and left Y-axis) and blood glucose levels (red line and right Y-axis) in mice treated with STZ at 200 mg/kg on Day 1 and Day 3 (n = 11). (**B**) Plasma insulin levels measured on Day 1 and Day 15 after STZ treatment. *N.D.* (not detected) indicates plasma insulin levels below the detection limit of the ELISA kit (MS303, Morinaga). n = 11. (**C**) Blood glucose levels after oral gavage of D-allulose at 1 g/kg in the STZ-induced type 1 diabetic mice (n = 10–11). Data are expressed as means ± SEM. Not significant by two-way ANOVA followed by Bonferroni’s test vs. saline group in **C**. og: oral gavage.

**Supplementary Fig. 3.**
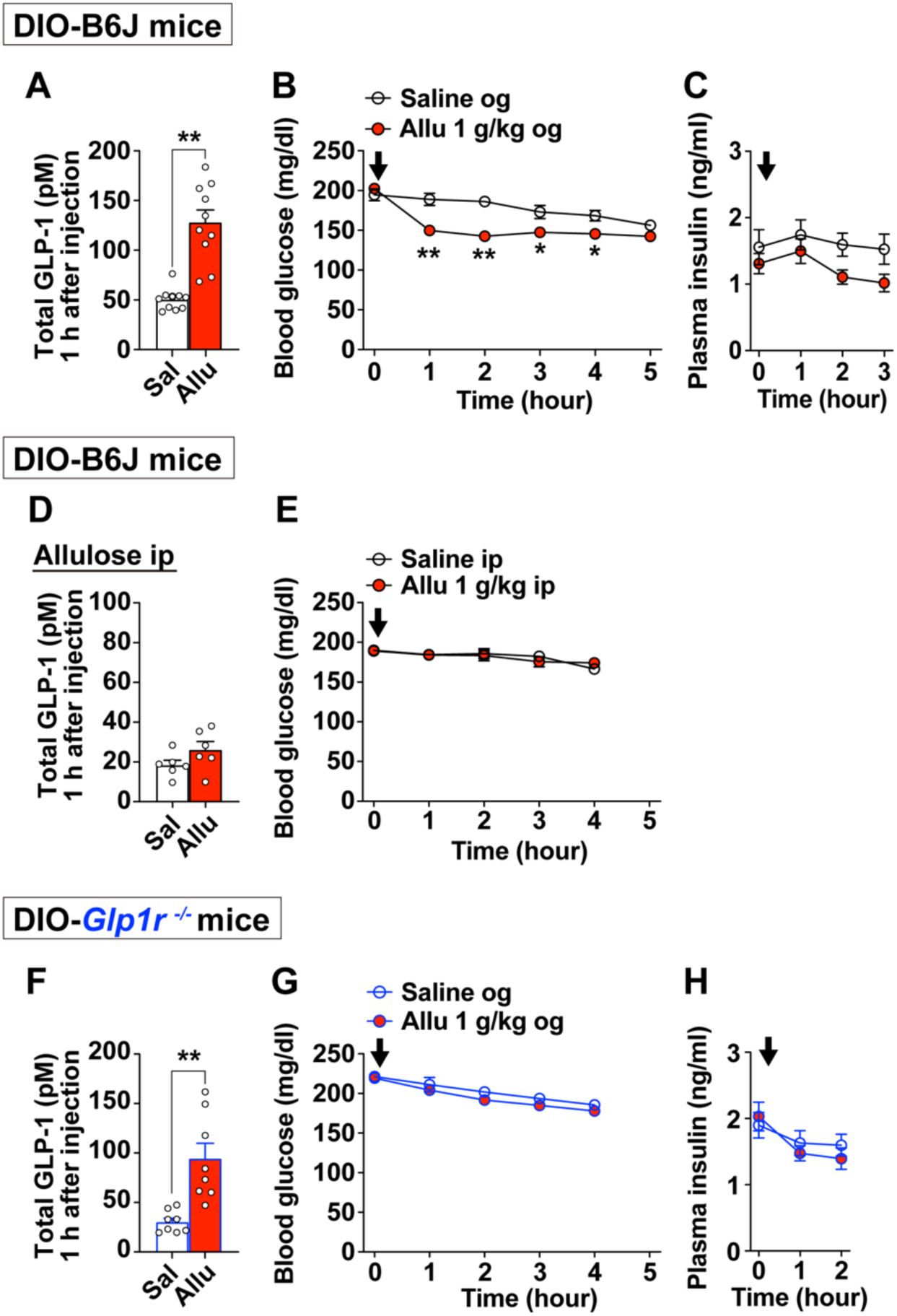
D-allulose-induced GLP-1 secretion improves hyperglycemia in diet-induced obese (DIO) mice via GLP-1 receptor signaling. (**A–E**) D-allulose at 1 g/kg or saline was administered orally (**A–C**) or intraperitoneally (**D, E**) to type 2 diabetic high-fat diet–induced obese C57BL/6J mice (DIO-B6J) fasted for 4 hours (n = 7–10). Plasma total GLP-1 levels in the portal vein were measured 1 hour after administration (**A, D**), along with sequential blood glucose levels (**B, E**) and plasma insulin levels (**C**). (**F–H**) DIO-*Glp1r* knockout mice fed a high-fat diet for 2 months to induce type 2 diabetes were orally gavaged D-allulose at 1 g/kg. Plasma total GLP-1 levels in the portal vein were measured 1 hour after administration (**F**), along with sequential blood glucose levels (**G**) and plasma insulin levels (**H**). Data are expressed as means ± SEM. **p < 0.01 by unpaired *t*-test (**A, F**), two-way ANOVA followed by Bonferroni’s test vs. saline group (**B**). Arrows in the figures indicate the timing of administration of each reagent. og: oral gavage.

**Supplementary Fig. 4.**
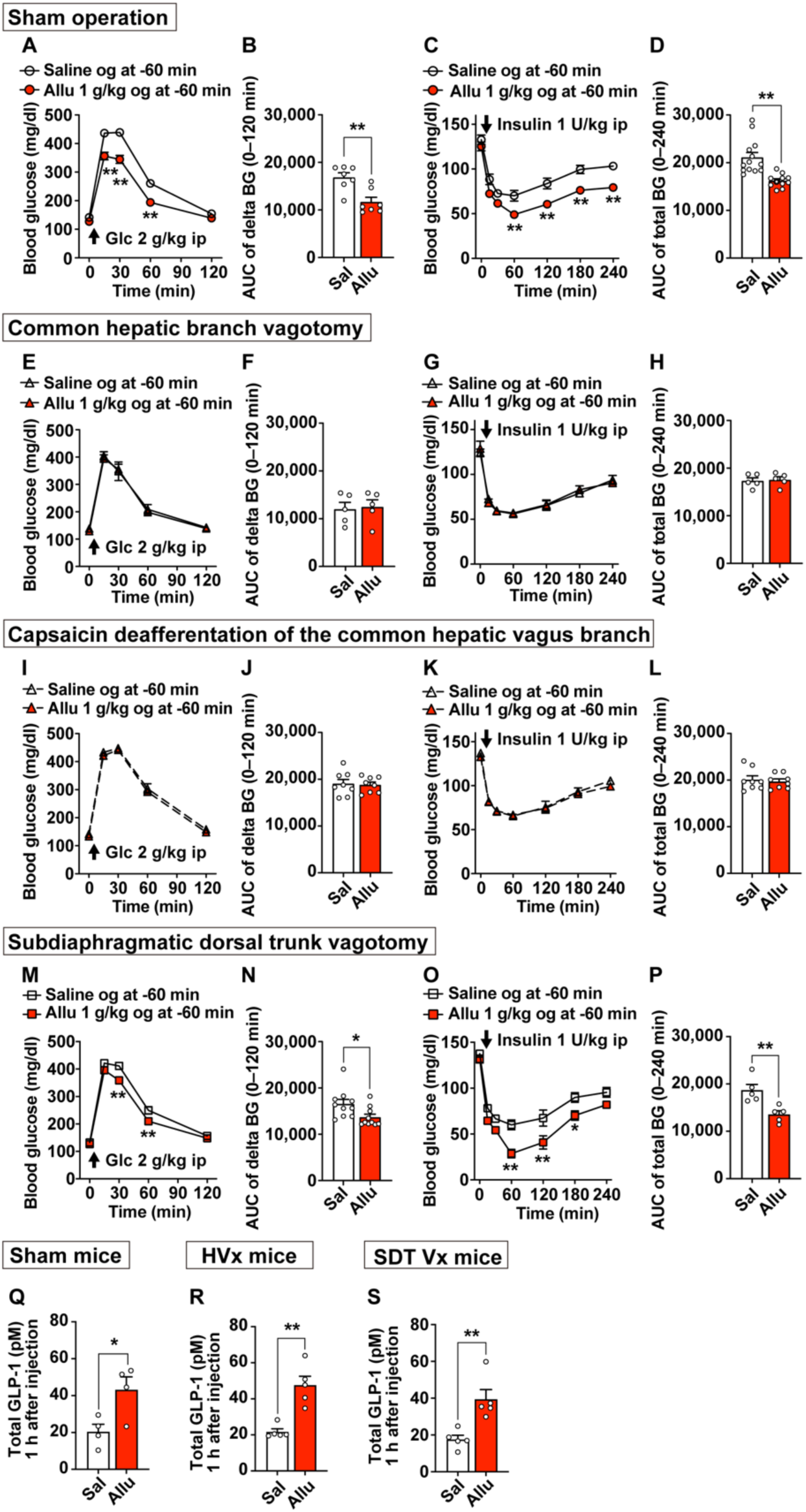
Left-sided, but not right-sided, vagus nerves are required for D-allulose-induced gut GLP-1 release to improve glucose tolerance and insulin sensitivity. (**A–D**) Four-hour fasted sham-operated C57BL/6J mice received oral gavage of saline or D-allulose at 1 g/kg 1 hour prior to ip injection of glucose (2 g/kg; **A**, **B**) or insulin (1 U/kg; **C**, **D**). n = 5–13. (**E–P**) The same experiments were performed in mice with common hepatic branch vagotomy (**E–H**, n = 5), capsaicin-induced deafferentation of the common hepatic branch of the vagus nerve (**I–L**, n = 8), and subdiaphragmatic dorsal trunk vagotomy (**M–P**, n = 5–10). (**Q–S**) Total GLP-1 levels in the portal vein plasma was measured 1 hour after oral gavage of saline or D-allulose at 1 g/kg in sham-operated mice (**Q**), hepatic vagotomized mice (**R**), and subdiaphragmatic dorsal trunk vagotomized mice (**S**). Data are expressed as means ± SEM. **p < 0.01, *p < 0.05 by unpaired *t*-test (**B, D, N, P, Q, R, S**), two-way ANOVA followed by Bonferroni’s test vs. saline group (**A, C, M, O**). Arrows in the figures indicate the timing of administration of each reagent. og: oral gavage.

**Supplementary Fig. 5.**
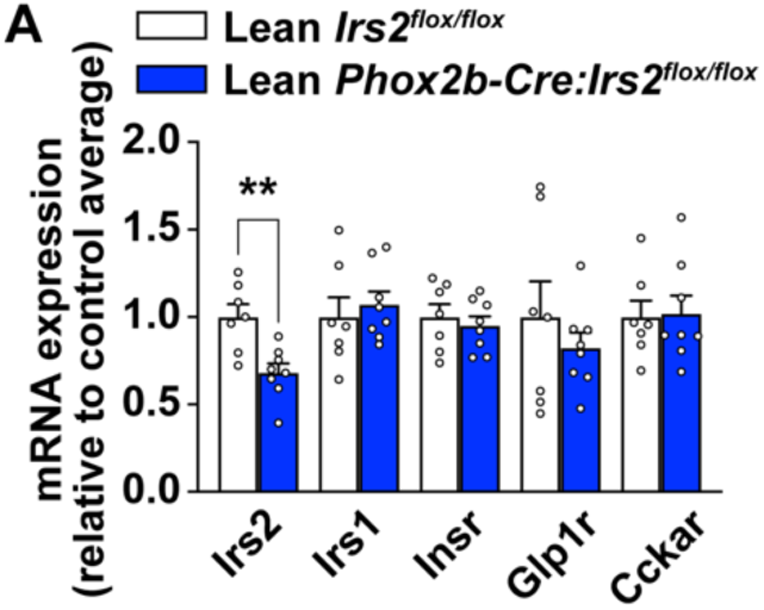
Expression of the hormone receptor mRNAs in the nodose ganglion of *Phox2b-Cre:Irs2^flox/flox^* mice. (**A**) Relative expression of *Irs2* and other hormone receptor mRNAs in the nodose ganglion in lean *Irs2^flox/flox^*and lean *Phox2b-Cre:Irs2^flox/flox^* mice fed standard chow (n = 7–8). Data are expressed as means ± SEM. **p < 0.01 by unpaired *t*-test.

**Supplementary Fig. 6.**
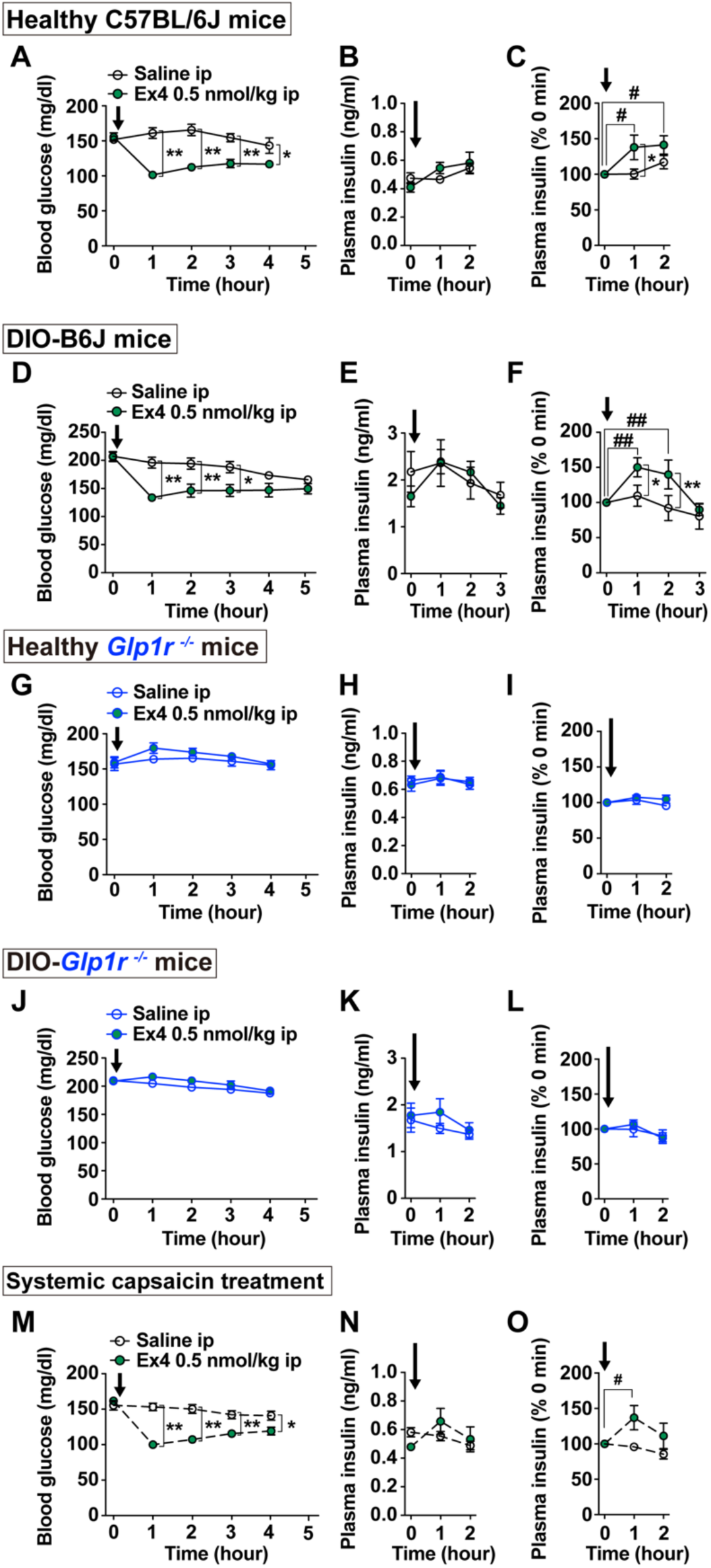
GLP-1 receptor agonist exendin-4 promotes insulin secretion and lowers blood glucose independently of sensory neural pathways. (**A–C**) Healthy C57BL/6J mice fasted for 4 hours received ip injection of GLP-1 receptor agonist exendin-4 (Ex4; 0.5 nmol/kg) or saline. Blood glucose levels (**A**) and absolute plasma insulin concentration (**B**) were measured, and relative plasma insulin levels (**C**) were calculated. n = 5. (**D–O**) Similar experiments were conducted in DIO-B6J mice fed high-fat diet (**D–F**, n = 6), lean *Glp1r* knockout mice fed standard chow (**G–I**, n = 6–7), DIO-*Glp1r* knockout mice fed high-fat diet (**J–L**, n = 8), and mice treated systemically with capsaicin to desensitize peripheral capsaicin-sensitive sensory neurons (**M–O**, n = 6–7). Data are expressed as means ± SEM. **p < 0.01, *p < 0.05 by two-way ANOVA followed by Bonferroni’s test vs. saline group (**A, C, D, F, M**). ##p < 0.01, #p < 0.05 by two-way ANOVA followed by Dunnett’s test vs. 0 h of each group (**C, F, O**). Arrows in the figures indicate the timing of administration of each reagent.

